# Three bacterial DedA subfamilies with distinct functions and phylogenetic distribution

**DOI:** 10.1101/2023.01.04.522824

**Authors:** Horia Todor, Nadia Herrera, Carol Gross

## Abstract

Recent studies in bacteria suggested that the broadly conserved but enigmatic DedA proteins function as undecaprenyl-phosphate (UndP) flippases, recycling this essential lipid carrier. To determine whether all DedA proteins have UndP flippase activity, we performed a phylogenetic analysis and correlated it to previously published experimental results and predicted structures. We uncovered three major DedA subfamilies: one contains UndP flippases, the second contains putative phospholipid flippases and is associated with aerobic metabolism, and the third is found only in specific Gram-negative phyla.

**IMPORTANCE:** DedA-family proteins are highly conserved and nearly ubiquitous integral membrane proteins found in Archaea, Bacteria, and Eukaryotes. Recent work revealed that eukaryotic DedA proteins are phospholipid scramblases and some bacterial DedA proteins are undecaprenyl phosphate flippases. We perform a phylogenetic analysis of this protein family in Bacteria revealing 3 DedA subfamilies with distinct phylogenetic distributions, genomic contexts, and putative functions. Our analysis lays the groundwork for a deeper understanding of DedA proteins and their role in maintaining and modifying the membrane.

## INTRODUCTION

DedA-family proteins are broadly distributed and nearly ubiquitous in eukaryotes, archaea, and bacteria. The structure of the DedA domain has not been solved, but computational (1) and topological (2) approaches indicate similarity to other transporter families (2). Eukaryotic VMP1, TVP38, and TMEM41B DedA proteins are central players in autophagy, while bacterial DedA family members play roles in colistin resistance, cell division, and pH sensitivity (detailed below).

Our understanding of DedA proteins was greatly bolstered by recent studies demonstrating that eukaryotic DedA homologs function as phospholipid scramblases (3–6) and that some bacterial DedA proteins are undecaprenyl phosphate (UndP) flippases (7, 8). UndP is the essential lipid carrier for the biogenesis of peptidoglycan and other bacterial surface polymers and must be recycled from the outer leaflet of the inner membrane to the cytoplasmic side – an essential function not previously associated with a gene.

These observations raise an important question: are all bacterial DedAs UndP flippases? Several factors suggest the answer is no. First, most bacteria have numerous DedAs: *Escherichia coli* has 8 and *B. subtilis* has 6. Second, only some DedAs exhibit UndP flippase activity (7). In *B. subtilis*, deletion of DedA homologs *yngC* and *ykoX* sensitized cells to the UndP-targeting drug MX2401, but deletion of the other 4 DedAs did not (7). In *E. coli* at least one DedA protein is required for viability (due to the essentiality of UndP flippase activity), but only 4/8 DedAs members are able to support viability as the sole DedA (9). Finally, some bacterial DedAs resemble eukaryotic DedAs, which are phospholipid scramblases (3–6).

To better understand the role(s) of bacterial DedAs, we performed a phylogenetic analysis of the DedA family in bacteria and found three major subfamilies, which largely correspond to the clusters of orthologous genes (COG) families COG0586, COG0398, and COG1238. Each subfamily exhibited distinct phylogenetic distribution, genomic context, and functional residues, implying distinct functions.

## RESULTS AND DISCUSSION

### DedA proteins are divided into 3 major subfamilies

To determine whether there are distinct DedA subfamilies within bacteria, we first identified all DedA homologs in ∼6,000 representative bacterial genomes using the PF09335 (DedA-family) motif (Materials and Methods, Table S1). We then used a representative subsample of these sequences to construct a tree (Materials and Methods, Fig. 1) and found three major subfamilies. These subfamilies corresponded to previously computed clusters COG0586, COG0398, and COG1238 and each contains DedAs from multiple bacterial phyla (Table 1, Fig. 1), suggesting that their divergence predated the last bacterial common ancestor.

**Table 1.** The distribution of DedA family proteins in different bacterial clades. Taxonomic assignment is based on the annotation level of the genome in eggNOG 4.5(40).

|  | COG0586 |  |  |  |  |  |  |  | COG0398 |  |  |  |  |  |  |  | COG1238 |  |  |  | Genomes |
| --- | --- | --- | --- | --- | --- | --- | --- | --- | --- | --- | --- | --- | --- | --- | --- | --- | --- | --- | --- | --- | --- |
|  | 0 | 1 | 2 | 3 | 4 | 5 | 6 | 7+ | 0 | 1 | 2 | 3 | 4 | 5 | 6 | 7+ | 0 | 1 | 2 | 3 |  |
| **γ-Proteobacteria** | 202 | 246 | 150 | 148 | 68 | 118 | 29 | 6 | 542 | 242 | 125 | 43 | 12 | 0 | 3 | 0 | 242 | 575 | 120 | 30 | **967** |
| **Actinobacteria** | 12 | 129 | 193 | 136 | 92 | 93 | 67 | 88 | 363 | 371 | 70 | 5 | 1 | 0 | 0 | 0 | 753 | 51 | 6 | 0 | **810** |
| **α-Proteobacteria** | 270 | 188 | 116 | 75 | 14 | 3 | 0 | 0 | 311 | 224 | 117 | 11 | 3 | 0 | 0 | 0 | 182 | 363 | 108 | 13 | **666** |
| **Bacilli** | 180 | 140 | 144 | 81 | 55 | 33 | 17 | 11 | 83 | 268 | 147 | 100 | 42 | 17 | 3 | 1 | 620 | 32 | 9 | 0 | **661** |
| **Clostridia** | 264 | 84 | 53 | 28 | 5 | 3 | 3 | 1 | 174 | 134 | 72 | 34 | 18 | 5 | 1 | 3 | 426 | 15 | 0 | 0 | **441** |
| **β-Proteobacteria** | 18 | 69 | 128 | 46 | 23 | 10 | 8 | 5 | 221 | 73 | 12 | 1 | 0 | 0 | 0 | 0 | 103 | 194 | 9 | 1 | **307** |
| **Flavobacteriia** | 106 | 73 | 4 | 4 | 0 | 0 | 0 | 0 | 163 | 20 | 3 | 1 | 0 | 0 | 0 | 0 | 143 | 44 | 0 | 0 | **187** |
| **Bacteroidia** | 25 | 126 | 20 | 0 | 0 | 0 | 0 | 0 | 171 | 0 | 0 | 0 | 0 | 0 | 0 | 0 | 28 | 137 | 6 | 0 | **171** |
| **Cyanobacteria** | 17 | 72 | 50 | 15 | 1 | 0 | 0 | 0 | 12 | 32 | 70 | 29 | 10 | 1 | 1 | 0 | 152 | 3 | 0 | 0 | **155** |
| **δ-Proteobacteria** | 35 | 37 | 27 | 13 | 4 | 2 | 0 | 0 | 26 | 30 | 31 | 25 | 5 | 1 | 0 | 0 | 58 | 48 | 12 | 0 | **118** |
| **ε-Proteobacteria** | 1 | 0 | 22 | 58 | 7 | 1 | 1 | 0 | 77 | 5 | 8 | 0 | 0 | 0 | 0 | 0 | 50 | 40 | 0 | 0 | **90** |
| **Tenericutes** | 88 | 0 | 0 | 0 | 0 | 0 | 0 | 0 | 85 | 3 | 0 | 0 | 0 | 0 | 0 | 0 | 86 | 2 | 0 | 0 | **88** |
| **Spirochaetes** | 33 | 29 | 20 | 0 | 0 | 0 | 0 | 0 | 69 | 10 | 2 | 1 | 0 | 0 | 0 | 0 | 60 | 22 | 0 | 0 | **82** |
| **Negativicutes** | 4 | 56 | 1 | 1 | 0 | 0 | 0 | 0 | 1 | 43 | 11 | 5 | 1 | 1 | 0 | 0 | 55 | 4 | 2 | 1 | **62** |
| **Cytophagia** | 26 | 9 | 15 | 2 | 2 | 0 | 0 | 0 | 6 | 38 | 9 | 1 | 0 | 0 | 0 | 0 | 48 | 6 | 0 | 0 | **54** |
| **Deinococcustherm** | 2 | 25 | 7 | 3 | 4 | 0 | 0 | 0 | 26 | 13 | 1 | 1 | 0 | 0 | 0 | 0 | 41 | 0 | 0 | 0 | **41** |
| **Sphingobacteriia** | 4 | 10 | 15 | 4 | 1 | 0 | 0 | 0 | 30 | 3 | 0 | 1 | 0 | 0 | 0 | 0 | 33 | 1 | 0 | 0 | **34** |
| **Fusobacteria** | 27 | 4 | 0 | 0 | 0 | 0 | 0 | 0 | 29 | 2 | 0 | 0 | 0 | 0 | 0 | 0 | 30 | 1 | 0 | 0 | **31** |
| **Erysipelotrichi** | 19 | 4 | 0 | 1 | 0 | 0 | 0 | 0 | 10 | 8 | 4 | 1 | 1 | 0 | 0 | 0 | 18 | 6 | 0 | 0 | **24** |
| **Chlamydiae** | 14 | 6 | 1 | 0 | 0 | 0 | 0 | 0 | 20 | 1 | 0 | 0 | 0 | 0 | 0 | 0 | 21 | 0 | 0 | 0 | **21** |
| **Thermotogae** | 16 | 1 | 0 | 0 | 0 | 0 | 0 | 0 | 17 | 0 | 0 | 0 | 0 | 0 | 0 | 0 | 17 | 0 | 0 | 0 | **17** |
| **Aquificae** | 8 | 4 | 3 | 1 | 0 | 0 | 0 | 0 | 16 | 0 | 0 | 0 | 0 | 0 | 0 | 0 | 3 | 11 | 2 | 0 | **16** |
| **Bacteria** | 3 | 10 | 1 | 2 | 0 | 0 | 0 | 0 | 7 | 9 | 0 | 0 | 0 | 0 | 0 | 0 | 15 | 1 | 0 | 0 | **16** |
| **Planctomycetes** | 2 | 8 | 5 | 0 | 0 | 1 | 0 | 0 | 7 | 5 | 1 | 1 | 2 | 0 | 0 | 0 | 13 | 2 | 1 | 0 | **16** |
| **Synergistetes** | 1 | 13 | 2 | 0 | 0 | 0 | 0 | 0 | 11 | 4 | 1 | 0 | 0 | 0 | 0 | 0 | 16 | 0 | 0 | 0 | **16** |
| **Chlorobi** | 0 | 0 | 7 | 6 | 0 | 0 | 0 | 0 | 13 | 0 | 0 | 0 | 0 | 0 | 0 | 0 | 12 | 1 | 0 | 0 | **13** |
| **Acidobacteriia** | 0 | 0 | 1 | 7 | 4 | 0 | 0 | 0 | 11 | 1 | 0 | 0 | 0 | 0 | 0 | 0 | 0 | 8 | 4 | 0 | **12** |
| **Verrucomicrobia** | 2 | 5 | 3 | 0 | 0 | 0 | 0 | 0 | 4 | 3 | 2 | 1 | 0 | 0 | 0 | 0 | 10 | 0 | 0 | 0 | **10** |

**Fig. 1.**
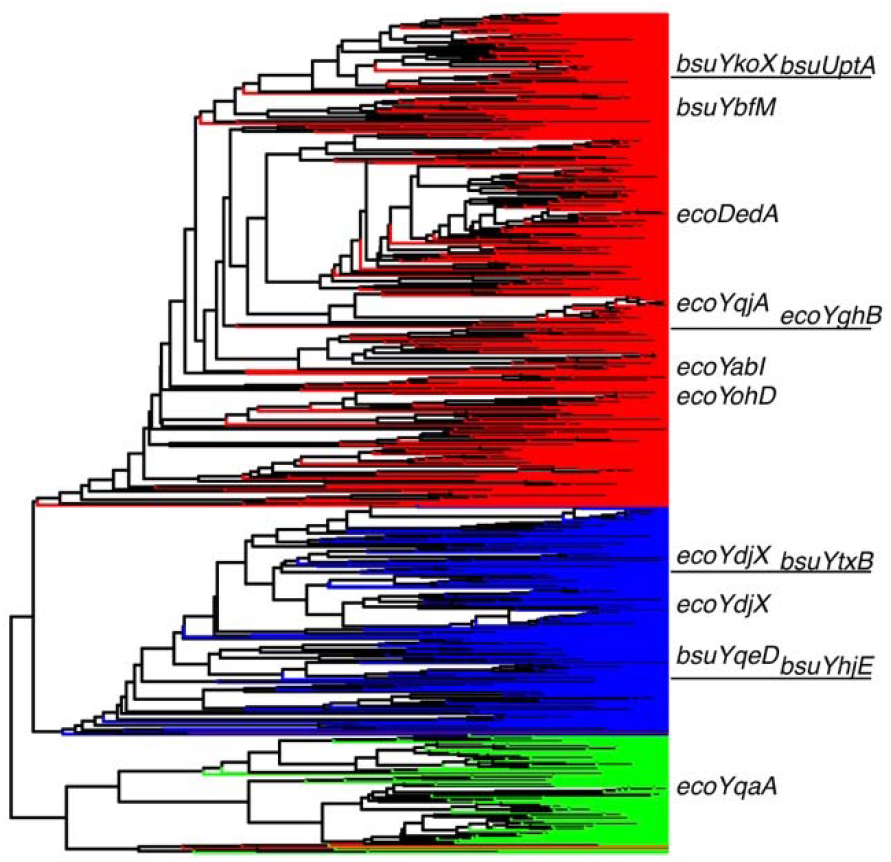
UPGMA tree of 1000 bacterial DedA proteins family reveals 3 distinct subfamilies, which are congruent with COG0586 (red), COG0398 (blue), and COG1238 (green). All DedA homologs from *E. coli* and *B. subtilis* are indicated.

We next performed an AlphaFold-based (10) structural analysis to assess differences between families (Fig. S1, Fig. 2C-E). We found that although the structure of the core DedA domain was largely conserved (RMSD 6-8Å), the placement of the N- and C-termini varied in both location and membrane orientation (cytoplasmic vs. periplasmic, Fig. S1). N- and C-terminal location and orientation were conserved within subfamilies (Fig. 2C-E), indicating that they are not caused solely by the mishandling of the C-terminal dimerization helices (2), but may reflect functional differences. The structural conservation of the DedA core domain and the variability of N- and C-terminal helices suggest that DedA subfamilies share a basic function (lipid scrambling/flipping) but vary in their substrate.

**Fig. 2.**
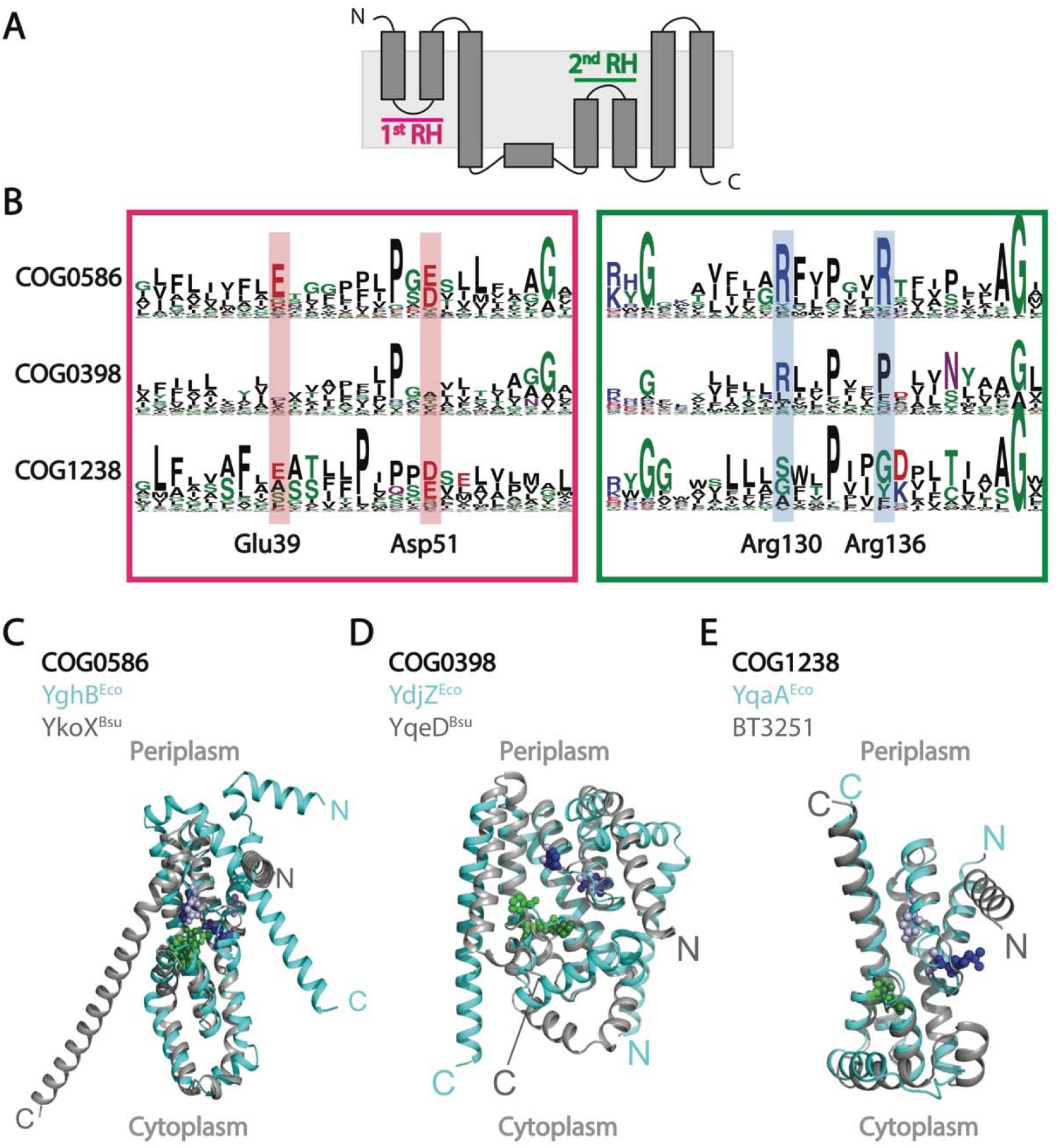
A) Schematic of core DedA domain topology from (2) depicting helices and re-entrant helices (RH). B) Consensus motif of the tips of the 1^st^ and 2^nd^ re-entrant helices in the 3 major DedA subfamilies, with conserved and essential residues highlighted. Residue numbering follows YghB^Eco^. C-E) AlphaFold models of representative DedA proteins in each family showing differing orientation of the N- and C-terminal helices and conserved structure of the core domain. Periplasmic and cytoplasmic orientation is based on topological study of YqjA^Eco^ (2). Colored spheres represent residues at the tip of the 1^st^ re-entrant helix (purple/light purple) and 2^nd^ re-entrant helix (green/dark green) for the first and second protein, respectively. RMSDs for the entire protein alignment and for the core DedA domain are 8.413Å/3.899Å for COG0586, 0.978Å/0.828Å for COG0398, and 0.968Å/0.972Å for COG1238.

### COG0586 DedA proteins are putative UndP flippases

The COG0586 subfamily accounts for ∼58% of bacterial DedAs. It is found in all major bacterial clades except for the peptidoglycan-less Tenericutes and the Thermotogota (Table 1). Almost all experimentally studied bacterial DedAs are in this family, including the sole (and essential) DedA of *Borrelia burgdoferi* (11), the target of the antibiotic halicyclamine A in *Mycobacterium smegmatis* (12), as well as DedAs implicated in colistin resistance through 4-amino-4-deoxy-L-arabinose modification of Lipid A in *Klebsiella pneumoniae* (13), *Enterobacter cloacae* (14), *Burkholderia thailandensis* (15), and *Burkholderia glumae* (16), and others (17–19).

Importantly, all DedA homologs thought to have UndP flippase activity (7, 8) are in this subfamily, including the four *E. coli* DedAs that can singly support viability (9) (and therefore putatively have UndP flippase activity), the 2 *P. aeruginosa* DedAs that can complement non-lethal phenotypes of a Δ2-DedA *E. coli* strain (20), and the 2 *B. subtilis* DedAs whose deletion affected MX2401 sensitivity (7). These data suggest that UndP flippase activity is the hallmark of the COG0586 DedA subfamily.

The conservation of key residues strengthens this hypothesis. The core DedA (1, 2) domain, which likely forms a homodimer (2), consists of an α-helical bundle with 2 re-entrant helices and 3 transmembrane helices (Fig. 2A): a conserved (Fig. S1) structure similar to other transporter families (2). In such transporters, the residues at the tips of the two re-entrant helices are invariably involved in substrate interactions. COG0586 DedAs contain conserved (Fig. 2) acidic residues at the tip of the 1^st^ re-entrant helix (YghB^Eco^ Asp51 and Glu39, Fig. 2B-C) that likely bind protons (2), and conserved (Fig. 2B-C) basic residues at the tip of the 2^nd^ re-entrant helix (YghB^Eco^ Arg130 and Arg136) that likely bind the negatively charged phosphate group of UndP (7). These residues were shown to be essential in both *E. coli* (21) and *B. subtills* (7) COG0586 proteins, and together likely effect PMF-driven flipping of UndP.

Genomic analyses provide additional evidence for the UndP flippase activity of the COG0586 subfamily. Although 93% of COG0586 family proteins contain only the PF09335 (DedA) domain, fusions to PF00581 (rhodanese) and PF01569 (PAP2) exist. PAP2 domains include phosphatidic acid phosphatases, which may dephosphorylate UndP-P prior to flipping; an essential function usually performed by a separate protein (e.g. BacA). Rhodanese domains are sulfurtransferases; their connection to UndP is unclear. Finally, COG0586 DedA genes are less common in genomes that also encode a non-DedA UndP flippase (7, 8) (COG2035, which contains DUF368) whereas no such relationship exists between COG2035 and the other subfamilies (Fig. S2). Together, these data suggest that COG0586 DedAs are UndP flippases, and raises the possibility that environmental or regulatory specialization is responsible for the genomic redundancy of COG0586 genes.

### COG0398 DedA proteins are associated with aerobic metabolism

The COG0398 DedA subfamily is the second largest family, accounting for ∼27% of bacterial DedA proteins. Intriguingly, eukaryotic DedAs such as TMEM41B, TMEM64, and VMP1 are most closely related to this bacterial subfamily (22).

COG0398 DedAs are unlikely to function as UndP flippases because they lack the acidic residues at the tip of the 1^st^ re-entrant helix associated with PMF-driven transport (Fig. 2B,D) and are missing one or both of the conserved basic residues at the tip of the 2^nd^ re-entrant helix putatively required for binding UndP (Fig. 2B,D) (7, 23). The sequence of the putative substrate binding region at the tip of the 2^nd^ re-entrant helix is similar to that of the eukaryotic DedA homologs, which have been characterized to function as phospholipid scramblases (3, 4). If bacterial COG0398 genes are also phospholipid scramblases, they may function to maintain or vary the phospholipid asymmetry in the inner membrane (24).

COG0398 DedAs exhibit a striking phylogenetic distribution (Table 1), being excluded from many predominantly anaerobic phyla (e.g. Bacteroidetes, Chlorobi) and classes (e.g. Bifidobacteriales, Propionibacteriales, Actinomycetales). Most COG0398 subfamily DedAs only contain the PF09335 (DedA) domain (90%). The most common domain fusions are with PF07992 and PF02852 domains (pyridine nucleotide-disulphide oxidoreductase & dimerization), which are found in glutathione/thioredoxin reductase, lipoamide dehydrogenase and mercuric reductase. Lipoamide and glutathione are associated with aerobic metabolism, consistent with the observed distribution of COG0398 DedAs. COG0398 DedAs proteins are essential in the α-proteobacterial model organisms *Caulobacter crescentus* and *Dinoroseobacter shibae* (25): further studies in these organisms may reveal their function and connection to aerobic metabolism.

### COG1238 subfamily DedA proteins are predominantly found in gram-negative bacteria

The COG1238 subfamily accounts for only ∼14% of DedA proteins: little is known about its function. A COG1238 homolog plays a role in indium resistance in *Rhodanobacter sp. B2A1Ga4*, but no mechanism was proposed (26).

COG1238 DedA subfamily proteins are also unlikely to function as UndP flippases; although the 1^st^ re-entrant helix exhibits conserved acidic residues suggestive of PMF-driven transport, the 2^nd^ re-entrant helix is almost completely lacking in positive residues that could bind a phosphate (Fig. 2B,E). The absence of conserved positive residues in the putative substrate binding site in the 2^nd^ re-entrant helix suggests that COG1238 subfamily DedAs may transport uncharged or even positively charged lipids. Almost all (99%) COG1238 subfamily proteins are single domain proteins.

The COG1238 subfamily DedAs are predominantly found in Proteobacteria, Bacteroidetes, Acidobacteriia, and Aquificae (>50% of species in these clades), occasionally in Cytophagia, Negativicutes, Planctomycetes, Flavobacteriia, Erysipelotrichi, and Spirochaetes (10-25%) and rarely in other gram-negative phyla or in gram-positive phyla such as Actinobacteria, Bacilli, and Clostridia (Table 1). It is unclear why this class is associated with such a specific subset of gram-negative bacteria. One possibility is that COG1238 DedAs transfer specific lipids to the inner leaflet of the outer membrane (OM), potentially through a direct interaction with an AsmA-family phospholipid bridge (27). Strikingly, this would mirror lipid transfers during autophagy, where eukaryotic DedA homologs transfer lipids to the autophagosome *via* ATG2, a protein homologous to bacterial AsmA proteins (28). COG1238 proteins are essential in several *Pseudomonas* species (25), opening the door to experimental characterization of their function.

### DedA proteins may be frequent antibiotic targets

The lipopeptide antibiotic amphomycin, which inhibits UndP recycling, was key to identifying the role of the COG0586 DedA-family (29, 30). We therefore asked whether additional antibiotics antagonize DedA activity. We reasoned that a modified DedA protein in the biosynthetic gene cluster (BGC) of such an antibiotic may provide immunity, as has been demonstrated for daptomycin (31) and speculated for others (31–33). We therefore searched the MIBiG database (34) of characterized BGCs for DedAs (PF09335) and identified 18 such clusters (Table S2). To more broadly ascertain DedA-containing BGCss, we next searched the antiSMASH database (35), which contains ∼147k computationally predicted BGCs, for clusters containing smCOG1188 (homologous to DedA) and identified 2213 DedA-containing BGCs in diverse bacteria including *Streptomyces, Bacillus*, and *Pseudomonas* (Table S2). Consistent with this broad distribution of putatively DedA-antagonizing antibiotics, oxydifficidin, a product of *Bacillus* species, was recently shown to kill *Neisseria gonorrhoeae* in a DedA dependent manner (36). These observations suggest that antagonizing DedA function is a widespread antibiotic modality.

## SUMMARY AND PERSPECTIVE

Recent experimental work in bacteria and eukaryotes has uncovered a conserved lipid flippase/scramblase function for DedA family proteins (3, 4, 6–8, 22). We show that bacterial DedA proteins share a conserved core structure (Fig. 2C-E, Fig. S1) but have evolved into three families with distinct functional residues and phylogenetic profiles (Fig. 1). These families likely predate the last bacterial common ancestor. The COG0586 family is likely involved in UndP recycling; the COG0398 family is associated with aerobic metabolism; and the COG1238 family is associated with the Gram-negative OM.

Studies of DedA family proteins in bacteria will shed new light on the diversity, function, and trafficking of bacterial lipids: topics which remain understudied. Understanding the substrate specificity, phenotypes, genetics, and regulation of the COG0398 DedA-family can elucidate the link between aerobic metabolism and the membrane, while understanding the COG1238 DedA-family can reveal novel aspects of the gram-negative outer membrane. Additionally, the putative presence of many DedA-antagonizing antibiotics in the genomes of *Actinobacteria* and *Bacilli* may provide useful membrane targeting antibiotics.

## ACKNOWLEDGMENTS

We thank T. Donohue, P. Kiley, P. Welander, and members of the C.A.G. laboratory for extensive helpful discussions, and L. Kim for help with Illustrator. N.H. was supported by the Burroughs Wellcome Fund Postdoctoral Enrichment Award #1019894, NIH T32 training grant (5T32HL007185-42) and the University of California President’s Postdoctoral Fellowship to NH. This work was supported by the National Institutes of Health grants R35 GM118061 (to C.A.G.).

## MATERIALS AND METHODS

### Identification of DedA homologs

To identify DedA homologs, we used hmmer 3.3.2 to query the proteomes of ∼6,000 representative bacterial genomes from the Progenomes1(37) database with the PF09335.14 motif characteristic of DedA-family proteins. This process identified and aligned ∼17,000 DedA homologs (Table S1). We excluded ∼600 proteins from poorly represented phyla and used the remaining set of 16,100 proteins for downstream analysis. The set of 16,100 proteins was annotated using eggNOG-mapper v2: 15,993 of the sequences were successfully annotated (>99%).

### Alignment

Because the PF09335.14 does not include the entirety of the 1^st^ re-entrant helix, we realigned the 16,100 sequences using the structure aware multiple-alignment program PROMALS3D(38).

### Tree construction

We constructed a tree using the DedA domain (PF09335.14) of 986 randomly chosen DedA proteins, plus all DedA homologs in *B. subtilis* and *E. coli* (a total of 1,000 sequences). Briefly, we considered only the columns of the alignment in which a least 95% of sequences had an aligned residue and calculated distances using BLOSUM62. The tree was constructed using the UPGMA algorithm and colored by protein membership in previously computed COGs (eggNOG 5.0(39, 40)).

### Structure prediction and analysis

Alphafold models were acquired from the alphafold database using the following identifiers: YghB^Eco^ (P0AA60), YdjZ^Eco^ (P76221), YqaA^Eco^ (P0ADR0), YkoX^Bsub^ (O34908), YqeD^Bsub^ (P54449), and BT3251 (Q8A2Q4). All models were aligned and modeled with PyMOL 2.5.4.

## SUPPLEMENTARY FIGURES

**Fig. S1.**
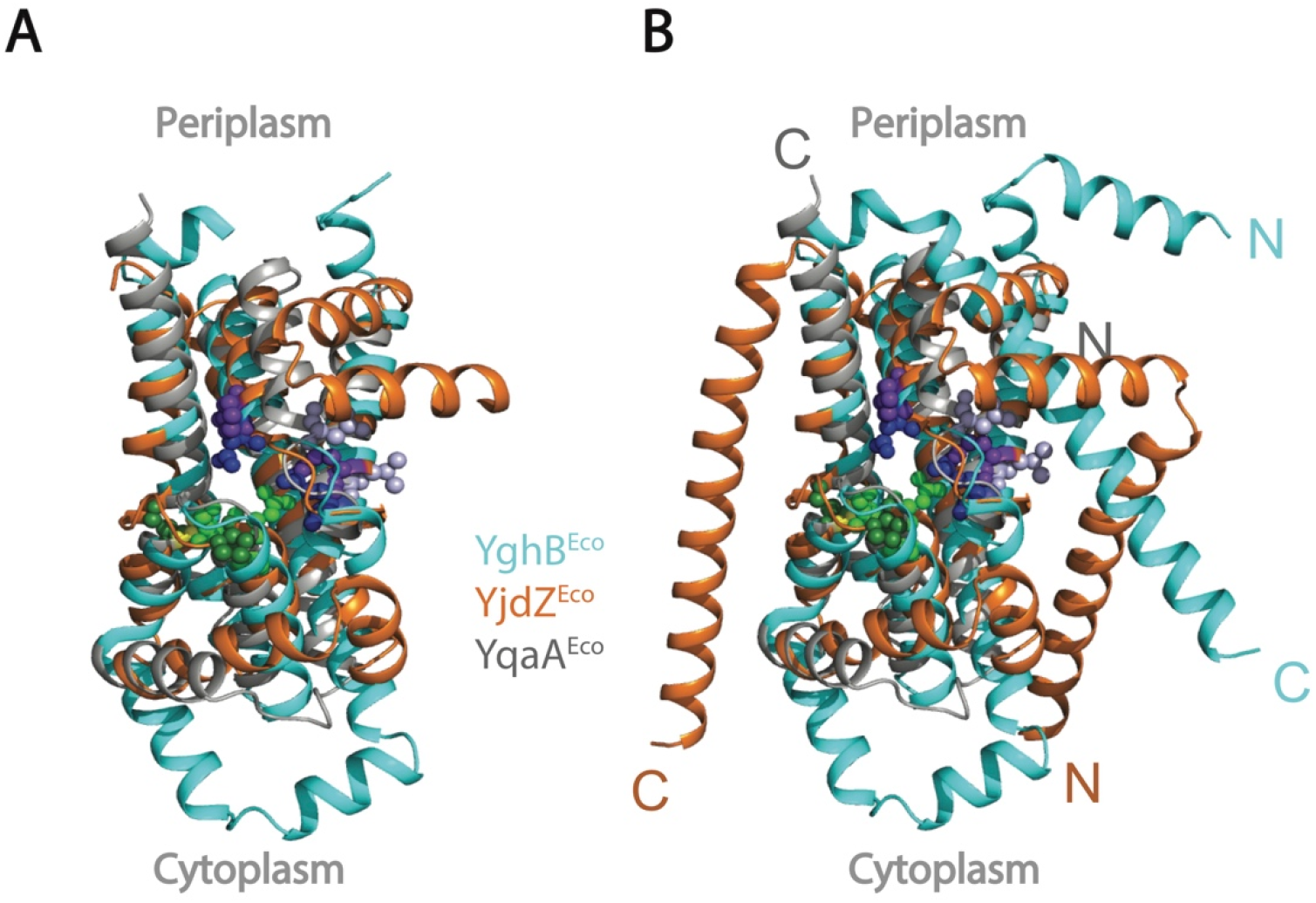
Aligned DedA from each of the three families from *E. coli*: YghB (cyan, COG0586), YdjZ (orange, COG0398), and YqaA (grey, COG1238). Alignment is shown as DedA domain only (A) or entire protein (B), highlighting the similarity of the core domain and the differences in the orientation of the N- and C-terminal helices. Periplasmic and cytoplasmic orientation is based on topological study of YqjA^Eco^ (2). RMSD of the DedA core domains: YghB-YqaA 8.579 Å, YghB-YdjZ 6.094 Å. RMSD of the entire protein: YghB-YqaA 8.578 Å, YghB-YdjZ 6.242 Å. Blue, purple, and light blue spheres show resides at the tip of the 1^st^ re-entrant helix for YghB, YqaA, and YdjZ, respectively. Green, yellow, and dark green spheres show resides at the tip of the 2^nd^ re-entrant helix for YghB, YqaA, and YjdZ, respectively.

**Fig. S2.**
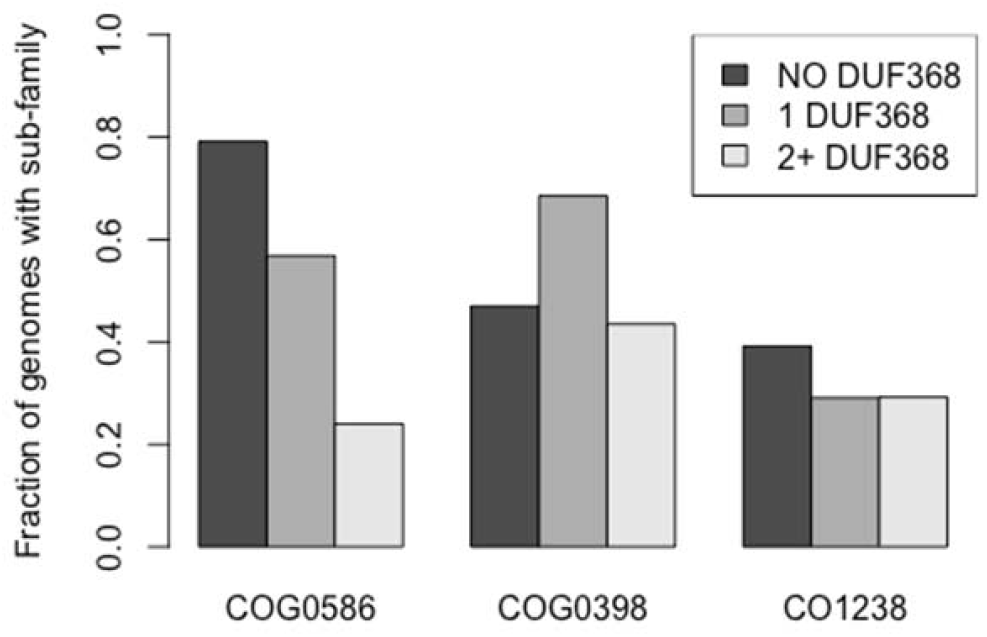
The proportion of genomes with a COG0586 family homolog is highest when no DUF368 family UndP flippases are present in the same genome, and decreases as the number of DUF368 members increase. No such pattern is observed for the other two subfamilies.

## SUPPLEMENTARY TABLES

**Table S1**. Contains information on all of the DedA homologs used in the analysis.

**Table S2**. Contains information about DedA homologs identified in BGCs.

